# Acetic acid activates distinct taste pathways in *Drosophila* to elicit opposing, state-dependent feeding responses

**DOI:** 10.1101/378414

**Authors:** Anita V. Devineni, Bei Sun, Anna Zhukovskaya, Richard Axel

## Abstract

Taste circuits are genetically determined to elicit an innate appetitive or aversive response, ensuring that animals consume nutritious foods and avoid the ingestion of toxins. We have examined the response of the fruit fly *Drosophila melanogaster* to acetic acid, a tastant that can be a metabolic resource but can also be toxic to the fly. Our data reveal that flies accommodate these conflicting attributes of acetic acid by virtue of a hunger-dependent switch in their behavioral response to this stimulus. Fed flies show taste aversion to acetic acid, likely a response to its potential toxicity, whereas starved flies show a robust appetitive response that may reflect their overriding need for calories. These opposing responses are mediated by two different classes of taste neurons. Acetic acid activates both the sugar and bitter pathways, which have opposing effects on feeding behavior. Hunger shifts the response from aversion to attraction by enhancing the appetitive sugar pathway as well as suppressing the aversive bitter pathway. Thus a single tastant can drive opposing behaviors by activating distinct taste pathways modulated by internal state.

## INTRODUCTION

Gustatory systems have evolved to identify appetitive substances of nutritional value and to elicit avoidance of toxic compounds. Organisms may encounter food sources that also contain harmful substances, and this poses an interesting perceptual problem. *Drosophila melanogaster,* for example, dines on fermenting fruit that contains both appetitive and aversive compounds. In anaerobic fermentation yeast and bacteria enzymatically convert six-carbon sugars into ethanol and acetic acid. These fermentation products can be toxic to the fly, yet the scent of decaying fruit is attractive and moreover the fly feeds despite the presence of these toxic compounds (McKenzie and Parsons, 1972; McKenzie and McKechnie, 1979; Zhu et al., 2003). These observations suggest that adaptive mechanisms may have evolved to ensure that toxic products do not impair the hungry fly from approaching and feeding on decaying fruits. Cider vinegar, for example, contains the potentially toxic metabolite acetic acid but elicits strong odor-evoked attraction (Semmelhack and Wang, 2009). *Drosophila melanogaster,* termed the vinegar fly, is more resistant to the toxic effects of acetic acid than other *Drosophila* species that do not depend upon fermenting food sources (McKenzie and McKechnie, 1979; Parsons, 1980). Moreover, flies can utilize acetic acid as a caloric source when deprived of other food sources (Parsons, 1980; Hoffmann and Parsons, 1984). Thus *D. melanogaster* may have evolved specific adaptations that allow the fly to recognize acetic acid as an appetitive tastant despite its potential toxicity. We have explored the seemingly paradoxical effects of acetic acid on feeding behavior in the vinegar fly.

Feeding is initiated by extension of the proboscis, a behavior that allows the fly to taste a potential food source (Dethier, 1976). Flies recognize a relatively small number of basic taste categories, including sweet, salty, bitter, fat, and carbonation (Liman et al., 2014; Zhang et al., 2013; Masek and Keene, 2013; Fischler et al., 2007). As in mammals, most tastants excite only one class of sensory neurons and each class is thought to activate determined neural pathways to elicit innate behavioral responses (Liman et al., 2014). For example, activation of sugar-responsive neurons drives appetitive feeding responses, whereas bitter-responsive neurons elicit aversion and suppress feeding (Liman et al., 2014; Marella et al., 2006).

The activation of gustatory neurons in flies and mammals elicits innate behavioral responses, but these responses can be modulated by internal states such as satiety or hunger. Hunger elicits several adaptive changes in behavior: increased food-seeking and food consumption, enhanced locomotor activity, decreased sleep, and altered olfactory and taste sensitivity (Sternson et al., 2013; Itskov and Ribeiro, 2013; Pool and Scott, 2014; Yang et al., 2015). In flies, starvation increases sugar sensitivity, which promotes feeding, and decreases bitter sensitivity, which enhances acceptance of food sources that contain aversive tastants (Inagaki et al., 2012; Inagaki et al., 2014). Starved flies also show enhanced olfactory attraction to cider vinegar, which facilitates food search behavior (Root et al., 2011). These hunger-dependent changes in both olfactory and gustatory sensitivity result, at least in part, from alterations in sensory neuron activity (Inagaki et al., 2012; Inagaki et al., 2014; Root et al., 2011).

Acetic acid, a product of fruit fermentation, signals the presence of food preferred by the fly and may also serve as a caloric source (Parsons, 1980; Hoffmann and Parsons, 1984). Acetic acid, however, can be toxic and flies avoid residing on food containing acetic acid (Parsons, 1980; Joseph et al., 2009). We have examined the behavioral and neural responses to the taste of acetic acid and observe that hunger induces a dramatic switch in the behavioral response to this metabolite. Fed flies show taste aversion to acetic acid whereas starved flies exhibit a strong appetitive response. Genetic silencing demonstrates that the bitter-sensing neurons mediate acetic acid aversion whereas the sugar-sensing neurons mediate the appetitive response to acetic acid. Hunger shifts the response from aversion to attraction by enhancing the sugar pathway as well as suppressing the bitter pathway. Two-photon calcium imaging reveals that acetic acid activates both sugar- and bitter-sensing neurons in both the fed and starved state. Thus, a single tastant activates two distinct neural pathways that elicit opposing behaviors dependent upon internal state. This hunger-dependent switch may reflect an adaptive response to acetic acid, a potential toxin that can also afford nutritional value under extreme conditions.

## RESULTS

### Acetic acid can elicit an appetitive or aversive taste response

The taste response to acetic acid was analyzed by examining the proboscis extension response (PER), an appetitive response that initiates feeding (Dethier, 1976). Appetitive tastants elicit PER when applied to the legs or labellum, the distal segment of the proboscis. Aversive tastants do not elicit PER and diminish the PER elicited by an attractive tastant. In fed flies, exposure of acetic acid (1-10%) to the labellum did not elicit PER (Figure 1A), even though these flies showed PER to sucrose (Figure 1B). Moreover, when acetic acid was mixed with sucrose it reduced the strong sugar-evoked PER to near-baseline levels (Figure 1C-D). 61% of fed flies exhibited PER to sucrose alone, and this response was reduced to 25% when 10% acetic acid was added (Figure 1C), a value close to that observed with water alone (Figure 1A-B). These data demonstrate that acetic acid elicits taste aversion in fed flies.

**Figure 1:**
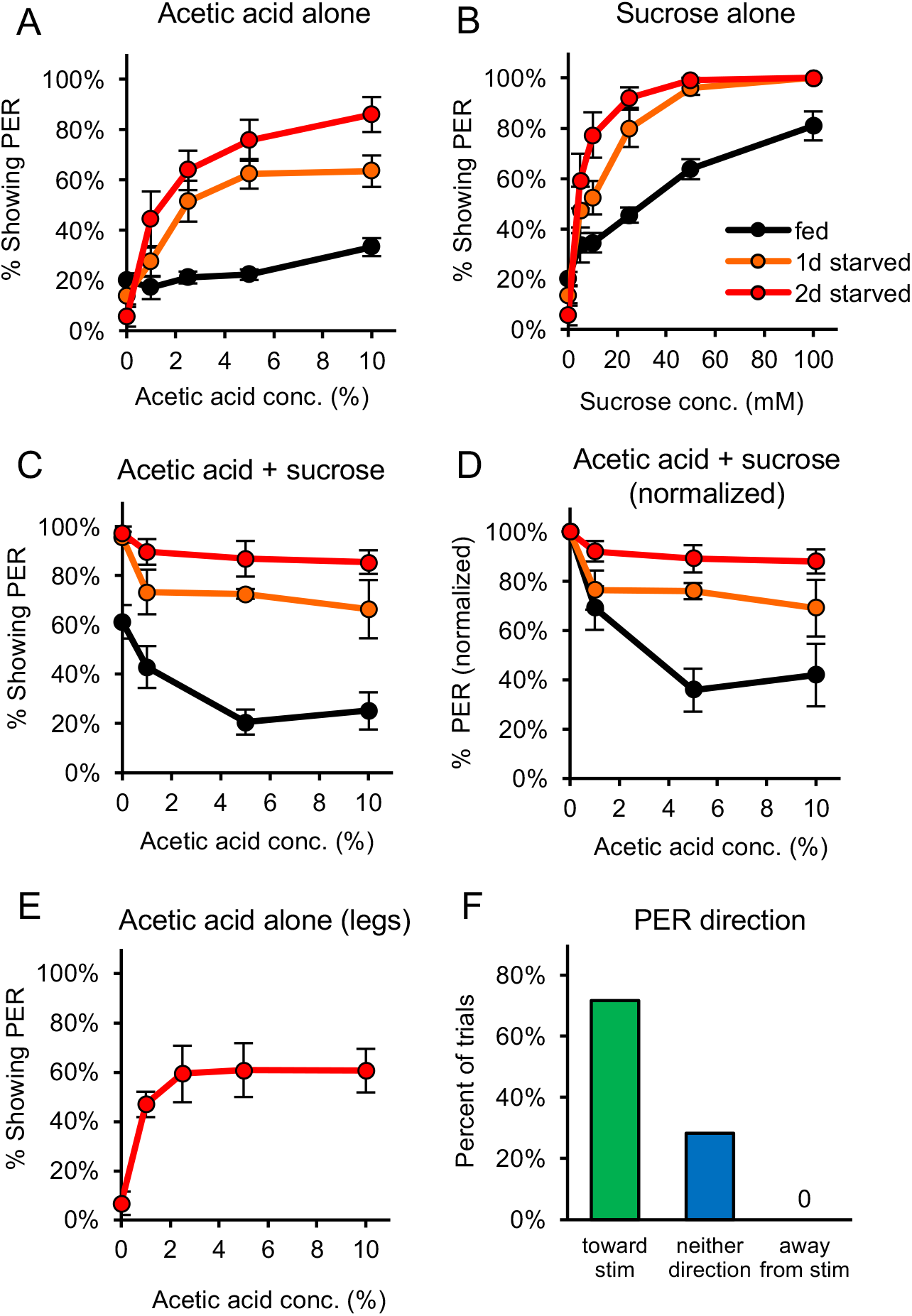
Acetic acid induces aversive or appetitive taste responses depending on hunger state. (A) One-day or two-day starved flies, but not fed flies, showed strong PER to acetic acid applied to the labellum. (B) Both fed and starved flies showed dose-dependent PER to sucrose applied to the labellum. (C-D) Acetic acid suppressed PER to sucrose in fed flies (p<0.01 at 5% and 10%), but not starved flies (p>0.05; one-way repeated-measures ANOVA followed by Dunnett’s post-tests for fed group comparing acetic acid to water). Panel D shows the same data in panel C normalized to the value for PER to 0% acetic acid in order to more easily compare groups. Acetic acid was mixed with 50 mM sucrose for starved flies and with 300 mM sucrose for fed flies to induce sufficient PER to observe potential suppression. (E) Two-day starved flies showed PER to acetic acid applied to the legs. (F) Two-day starved flies stimulated asymmetrically with acetic acid on the legs tended to show PER in the direction toward the stimulus (n = 53 trials, 9 flies).

A dramatic switch was observed in the behavioral response of starved flies. When acetic acid was applied to the labellum of starved flies, strong, dose-dependent PER was observed, with 86% of flies exhibiting PER to 10% acetic acid after two days of starvation (Figure 1A). Moreover, when acetic acid was added to sucrose no suppression of sugar-evoked PER was observed in starved flies, a finding in sharp contrast to the PER suppression in fed flies (Figure 1C-D). This switch from an aversive response in fed flies to an appetitive response in starved flies is specific for acetic acid and was not observed for other aversive tastants, such as the bitter compounds quinine and lobeline (Figure 1 – figure supplement 1). Appetitive compounds such as sugar also do not elicit a qualitative switch in behavior, since sugar elicited PER in both fed and starved flies (Figure 1B). Thus a single tastant, acetic acid, can elicit opposing behavioral responses dependent upon internal state.

We next performed experiments to demonstrate that PER elicited by acetic acid is a component of an appetitive feeding response. When a fly is stimulated asymmetrically with an appetitive tastant on only one leg, extension of the proboscis is observed in the direction of the stimulus (Saraswati, 1998; Schwarz et al., 2017). We first confirmed that acetic acid elicits strong PER in starved flies when applied to the legs instead of the labellum (Figure 1E). We then stimulated the legs asymmetrically and observed that in 72% of trials, starved flies that showed PER extended the proboscis in the direction of the stimulus (Figure 1F, Video 1). Proboscis extension was never observed in the direction opposing the stimulus, and in 28% of trials flies exhibited PER neither toward nor away from the stimulus (Figure 1F). We also observed that when afforded the option to consume 5% acetic acid following PER, 7 of the 10 flies tested consumed it (Video 2). Thus, the majority of flies extend their proboscis in the direction of an acetic acid stimulus and voluntarily consume it, suggesting that this response is an appetitive component of feeding behavior.

Acetic acid exists in solution as three chemical species: undissociated acetic acid, which partially dissociates to produce acetate and protons. We asked whether PER to acetic acid reflects a more general taste response to low pH. Starved flies failed to show PER to hydrochloric acid at pH values equivalent to those of 5% or 10% acetic acid, which elicit strong PER, indicating that low pH is not sufficient to induce an appetitive response (Figure 1 – figure supplement 2A). We also tested the response of starved flies to potassium acetate at molarities equivalent to those of 5% or 10% acetic acid and failed to observe a response (Figure 1 – figure supplement 2B). These experiments suggest that neither protons nor acetate ions are capable of eliciting PER, suggesting that undissociated acetic acid is recognized by taste cells. In accord with this suggestion, propionic acid, a simple carboxylic acid structurally similar to acetic acid, elicited strong PER in starved flies whereas the more distantly related citric acid elicited a weaker response (Figure 1 – figure supplement 2C). Thus, undissociated small aliphatic acids may be recognized by gustatory neurons to elicit PER in starved flies.

### PER to acetic acid is mediated by the gustatory system

Proboscis extension can be elicited by the taste organs, but it remains possible that other sensory modalities such as olfaction contribute to this behavioral response. Acetic acid activates olfactory sensory neurons (Ai et al., 2010). We therefore removed the olfactory organs, the third antennal segment and maxillary palp, from two-day starved flies and observed that PER to acetic acid was unperturbed by this manipulation (Figure 2A). 76% of flies lacking olfactory organs showed PER to 10% acetic acid, a value close to that observed with control flies (72%; Figure 2A). Fed flies lacking olfactory organs failed to show PER to acetic acid, mirroring the behavior of control flies (Figure 2B). Fed flies lacking olfactory organs also continued to show aversion to acetic acid, as we observed significant suppression of PER when acetic acid was added to sucrose (Figure 2C). These experiments demonstrate that both the appetitive and aversive proboscis extension responses to acetic acid are observed in the absence of olfactory organs, and are likely to be mediated by the gustatory system.

**Figure 2:**
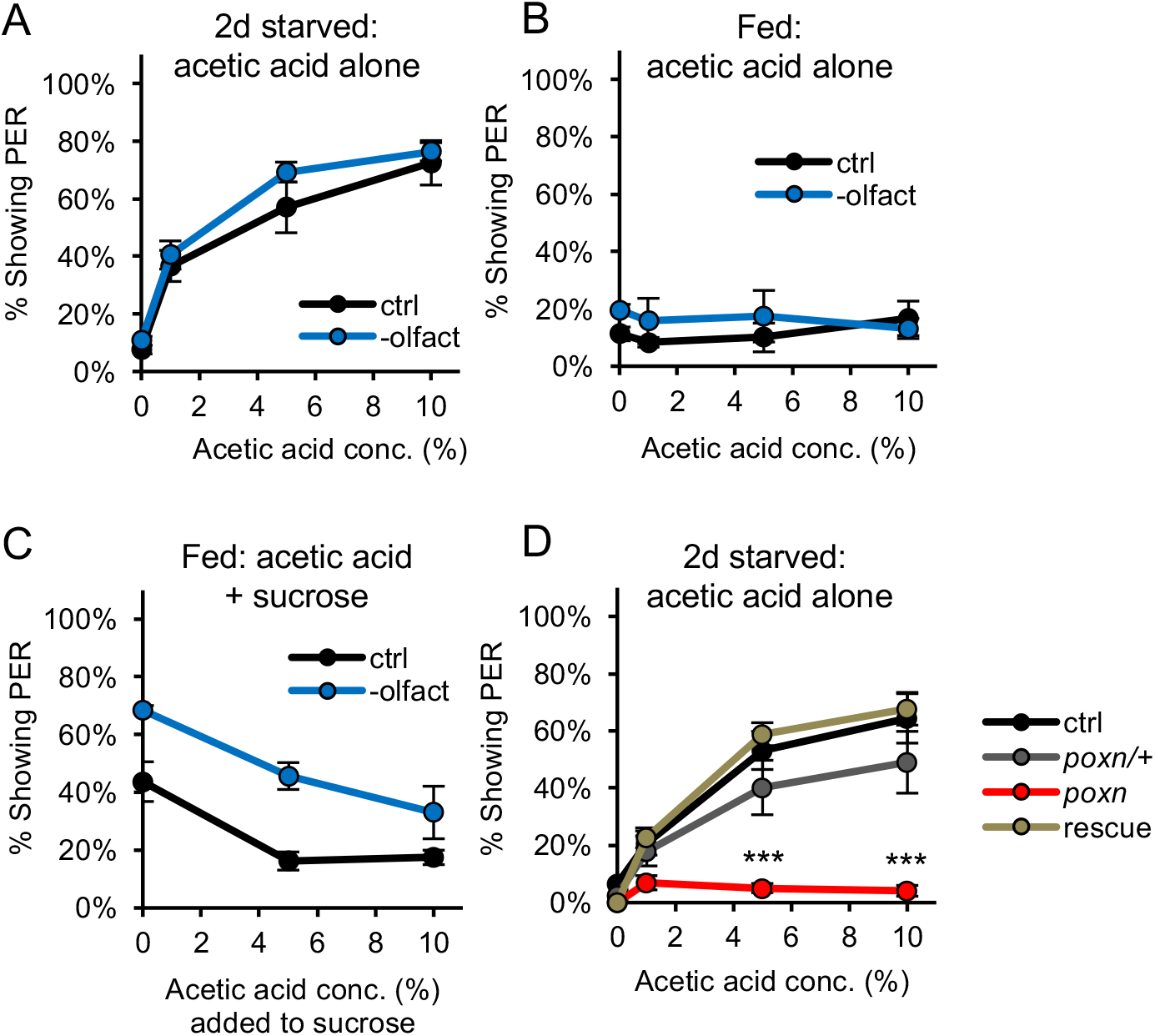
PER to acetic acid is mediated by the gustatory system, not the olfactory system. (A-B) Removing the olfactory organs did not affect PER to acetic acid in two-day starved flies (A) or fed flies (B) (p>0.05, two-way repeated measures ANOVA). (C) Acetic acid aversion in fed flies, measured by suppression of PER to 50 mM sucrose, was observed in both control flies (p<0.01) and flies lacking olfactory organs (p<0.05, one-way repeated measures ANOVA). Flies without olfaction were generally more responsive than control flies (p<0.01, two-way repeated measures ANOVA). (D) Two-day starved flies homozygous for the *poxn^ΔM22-B5^* mutation showed decreased PER to acetic acid as compared to wild-type controls, *poxn^ΔM22-B5^/+* heterozygotes, and *poxn^ΔM22-B5^* homozygotes carrying a rescue transgene (***p<0.001, two-way repeated measures ANOVA followed by Bonferroni post-tests).

We demonstrated the requirement for taste neurons by examining the response to acetic acid in *pox-neuro (poxn)* mutants, in which taste bristles are transformed into mechanosensory bristles lacking gustatory receptors (Boll and Noll, 2002). Starved *poxn^ΔM22-B5^* homozygous mutants failed to display PER to any concentration of acetic acid tested (Figure 2D). In contrast, wild-type flies, *poxn^ΔM22-B5^/+* heterozygotes (which have normal bristles), and rescue flies (*poxn^ΔM22-B5^* mutants carrying the *SuperA* rescue transgene) showed strong PER, with up to ~50-70% of flies responding (Figure 2D). Interpretation of these experiments must be tempered by the observation that the *poxn^ΔM22-B5^* mutants often appeared physically smaller than control flies and are known to have central nervous system abnormalities in addition to their lack of taste bristles (Boll and Noll, 2002). Nonetheless, these experiments suggest that the appetitive and aversive responses to acetic acid require the taste organs and are largely independent of olfaction.

### Sugar-sensing neurons mediate PER to acetic acid

Neurons in the chemosensory bristles of the labellum detect distinct taste modalities, including sugar, bitter, water, and low and high concentrations of salt (Liman et al., 2014; Marella et al., 2006; Cameron et al., 2010; Zhang et al., 2013). We employed genetic silencing to identify the neuronal classes responsible for the appetitive and aversive responses to acetic acid. Sugar-sensing neurons express multiple chemoreceptors and elicit PER in response to sugars (Dahanukar et al., 2007; Slone et al., 2007; Fujii et al., 2015). The receptor Gr64f is expressed in all sugar-responsive taste neurons (Fujii et al., 2015). We therefore silenced the sugar-sensing neurons by expressing *UAS-Kir2.1,* encoding an inwardly rectifying potassium channel (Baines et al., 2001), under the control of the transcriptional activator *Gr64f-Gal4.* Starved flies harboring both the *Gr64f-Gal4* and *UAS-Kir2.1* transgenes showed very low frequencies of PER (12%-28%) in response to increasing concentrations of either sucrose or acetic acid (Figure 3A-B). Control flies containing either the *Gr64f-Gal4* or *UAS-Kir2.1* transgenes alone resembled wild-type flies and exhibited strong PER to both sucrose and acetic acid, with up to 100% of flies responding to sucrose and up to ~70-80% responding to acetic acid (Figure 3A-B). These experiments demonstrate that the appetitive response to acetic acid observed in starved flies is mediated by the sugar-sensing neurons.

**Figure 3:**
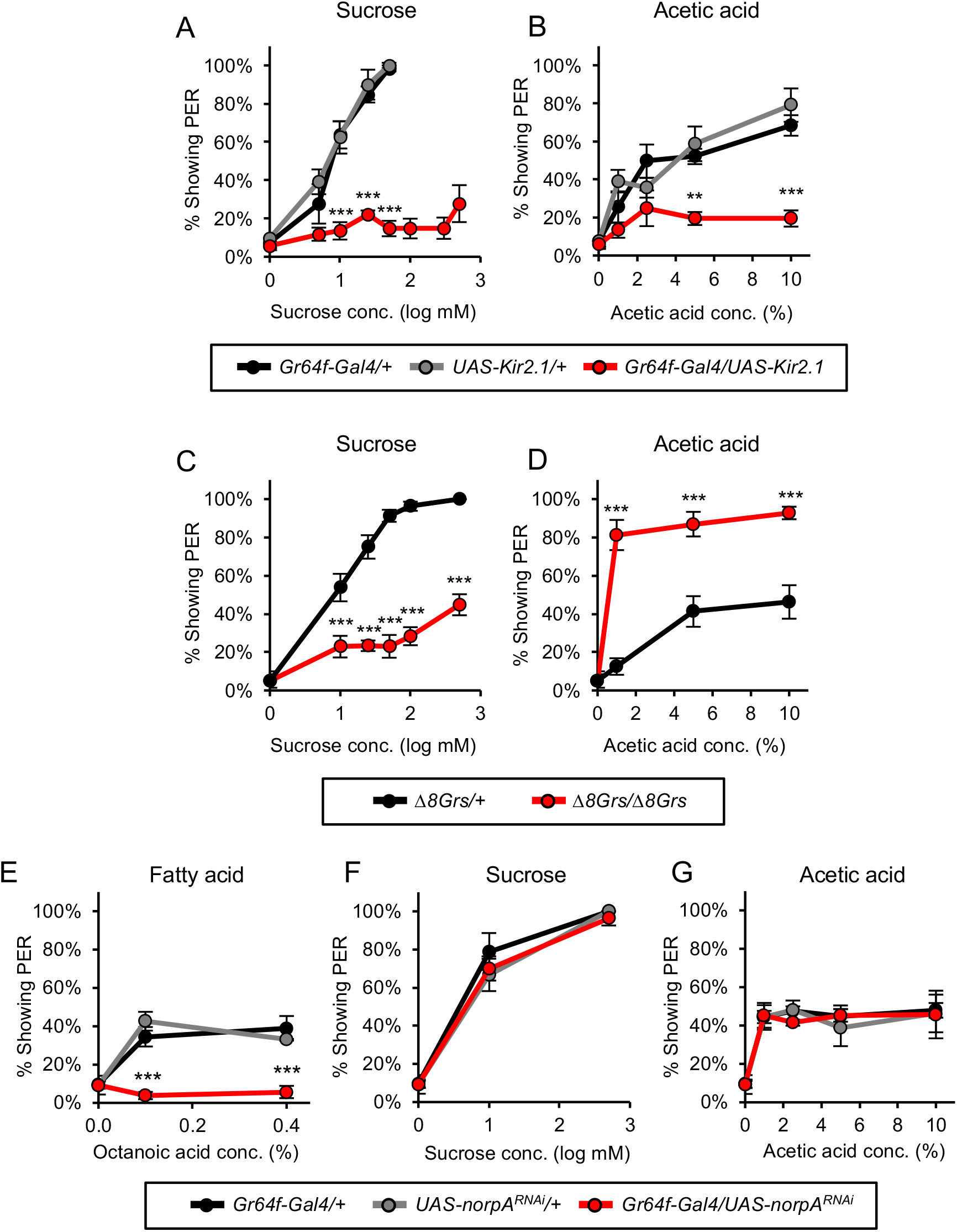
Sugar-sensing neurons mediate PER to acetic acid in starved flies. (A-B) Silencing the activity of sugar-sensing neurons impaired PER to sucrose (A) and acetic acid (B) in two-day starved flies. (C-D) One-day starved homozygous mutant flies lacking all eight sugar receptors showed decreased PER to sucrose (C) but showed increased PER to acetic acid (D) relative to heterozygous controls. (E-G) RNAi knockdown of *norpA,* encoding PLCβ, in sugar-sensing neurons abolished PER to fatty acid (E) but did not affect PER to sucrose (F) or acetic acid (G) in two-day starved flies. For all panels: **p<0.01, ***p<0.001, two-way repeated measures ANOVA followed by Bonferroni’s post-tests comparing experimental group to each control group. In panel A, statistical analyses did not include the three highest sucrose concentrations because control flies were not tested at these concentrations.

We asked whether the acetic acid response is mediated by the gustatory receptors (Grs) that detect sugars. Eight Grs are expressed in sugar sensory neurons (Fujii et al, 2015; Yavuz et al., 2015). As expected, homozygous flies carrying deletions in all eight sugar-sensing Gr genes *(ΔβGrs/ΔβGrs;* Yavuz et al., 2015) showed a strong reduction in PER to sucrose as compared with control heterozygous flies (Figure 3C). In contrast to their reduced response to sugar, homozygous mutant flies continued to show PER to acetic acid (Figure 3D). Interestingly, acetic acid-evoked PER in homozygous mutant flies was significantly greater than in control flies (Figure 3D). This increase in PER to acetic acid may reflect the possibility that hunger is intensified in mutants lacking sugar receptors and this may enhance the sensitivity of the sugar-sensing circuit. Alternatively, the acetic acid and sucrose transduction pathways may employ a common limiting component that is no longer limiting in mutant sugar-sensing neurons. These experiments demonstrate that the response to acetic acid in starved flies is mediated by the sugar-sensing neurons but does not employ the sugar receptors.

The sugar-sensing neurons also elicit PER in response to fatty acids, such as hexanoic and octanoic acid, through a molecular mechanism distinct from sugar detection (Masek and Keene, 2013; Ahn et al., 2017). We therefore asked whether sugar neurons recognize acetic acid and fatty acids through the same mechanism, since hexanoic and octanoic acids also are aliphatic carboxylic acids. A previous study showed that PER induced by fatty acids requires phospholipase C (PLC) signaling in sugar-sensing neurons whereas PLC is dispensable for PER to sucrose (Masek and Keene, 2013). We therefore tested whether PLC signaling in sugar neurons is required for PER to acetic acid. An RNAi transgene targeting the gene *norpA,* a fly ortholog of PLC, was expressed in sugar neurons under the control of *Gr64f-Gal4.* RNAi inhibition of *norpA* expression in the sugar neurons severely reduced PER to fatty acids whereas PER to either sucrose or acetic acid was unaffected (Figure 3E-G). These data suggest that the appetitive response to acetic acid is mediated by sugar-sensing neurons, but engages molecular pathways distinct from those employed in the detection of either sugars or fatty acids.

### Bitter-sensing neurons suppress PER to acetic acid

We next identified the neurons that mediate the aversive response to acetic acid in fed flies. Multiple classes of bitter sensory neurons reside in the labellum, and each bitter neuron expresses the receptor *Gr66a* (Weiss et al., 2011). We therefore employed the regulatory sequences of *Gr66a* to drive the expression of *Kir2.1* to silence the bitter neurons. In initial experiments we demonstrated the efficacy of Kir2.1 silencing. Starved control flies exhibit PER to sucrose, and this response is strongly diminished by the addition of the bitter compounds quinine or lobeline (Figure 4 – figure supplement 1A-B). This suppression of PER by bitter compounds is no longer observed when Kir2.1 is expressed in bitter neurons, demonstrating the efficacy of Kir2.1 silencing (Figure 4 – figure supplement 1A-B). We therefore employed Kir2.1 to examine the effect of silencing bitter neurons on the aversive responses to acetic acid in fed flies. In control fed flies acetic acid suppressed PER to sucrose, but this suppression was largely eliminated when the bitter neurons were silenced (Figure 4A). In controls, ~80% of flies exhibited PER to sucrose alone, and this was reduced to ~20-30% upon addition of 10% acetic acid. In contrast, when bitter neurons were silenced over 80% of flies continued to exhibit PER upon exposure to a mixture of sugar and 10% acetic acid (Figure 4A). These results indicate that bitter-sensing neurons mediate the aversive response to acetic acid in fed flies.

**Figure 4:**
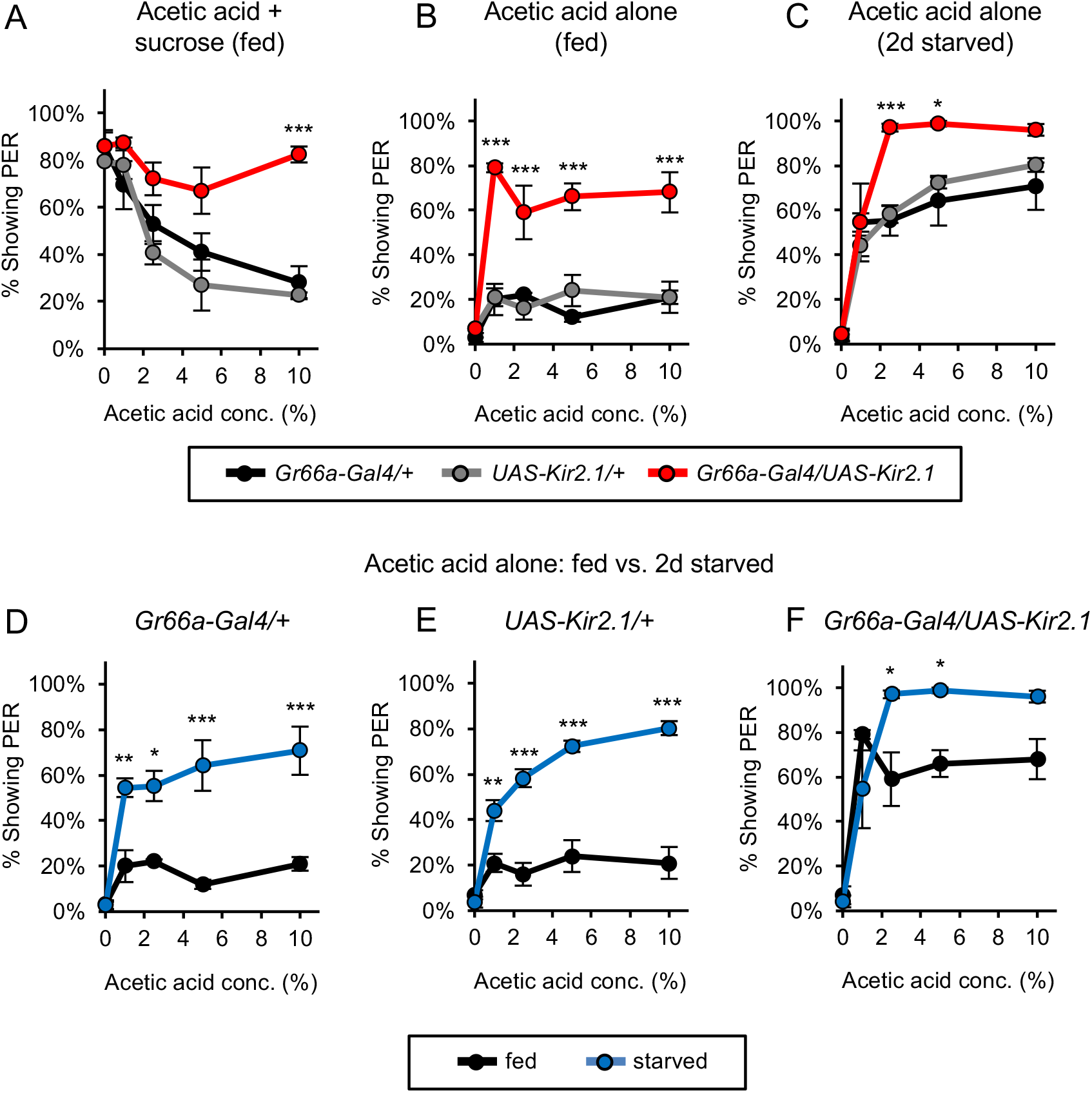
Bitter-sensing neurons suppress PER to acetic acid. (A) Silencing bitter-sensing neurons strongly reduced aversion to acetic acid in fed flies. Aversion was measured as the suppression of PER to 100 mM sucrose containing acetic acid. Both sets of control flies showed significant aversion to acetic acid (p<0.001), whereas experimental flies did not show significant aversion (p>0.05, one-way repeated measures ANOVA). (B-C) Silencing bitter-sensing neurons enhanced PER to acetic acid alone in fed flies (B) and two-day starved flies (C). (D-F) Comparing fed and starved flies of each genotype (same data as panels B and C) revealed that starvation enhanced PER in all genotypes. For all panels: *p<0.05, **p<0.01, ***p<0.001, two-way repeated measures ANOVA followed by Bonferroni’s post-tests comparing experimental group to each control group.

We have also examined the consequences of bitter neuron silencing on the responses to acetic acid alone. Starved flies exhibit PER to acetic acid alone, but this response is not observed in fed flies (Figure 1A). Acetic acid elicited PER in ~20% of control fed flies, a value near baseline (Figure 4B). Silencing of the bitter neurons resulted in a striking increase in the frequency of fed flies that exhibited PER to acetic acid (~60-80%; Figure 4B). Silencing bitter neurons in fed flies did not affect PER to sucrose alone, indicating that the bitter neurons do not exert nonspecific suppression of PER (Figure 4 – figure supplement 1C). These results afford an explanation for the observation that fed flies normally fail to exhibit PER to acetic acid. Our data suggest that acetic acid activates sugar-sensing neurons, which promote an appetitive response, but in the fed state simultaneous activation of bitter-sensing neurons completely suppresses this response. Silencing the bitter neurons eliminates this suppression, unmasking the appetitive response.

Silencing the bitter neurons resulted not only in the emergence of PER to acetic acid in fed flies but also enhanced PER to acetic acid in starved flies (Figure 4C). PER to 10% acetic acid was observed in 70-80% of control flies and this value increased to 96% upon bitter neuron silencing (Figure 4C). Silencing bitter neurons in starved flies did not affect PER to sucrose (Figure 4 – figure supplement 1D). These results demonstrate that even in the starved state, bitter neurons suppress PER to acetic acid. The observation that bitter neuron silencing has a stronger effect on acetic acid-induced PER in fed flies (Figure 4B) than in starved flies (Figure 4C) suggests that hunger inhibits the bitter pathway.

The striking increase in PER to acetic acid after starvation results from the hunger-dependent suppression of the bitter-sensing pathway but may also reflect enhancement of the appetitive sugar-sensing pathway. We and others observe that PER to sucrose is increased by starvation, indicating that the sugar pathway is upregulated by hunger (Figure 1B; Inagaki et al., 2012). We therefore examined the relative contributions of the sugar- and bitter-sensing pathways to acetic acid-induced PER. We compared responses to acetic acid in both fed and starved flies with and without bitter neuron silencing. In control flies, starvation strongly increased PER to acetic acid: 20% of fed flies and 70-80% of starved flies responded at the highest concentration (Figure 4D-E). Upon bitter neuron silencing the difference between PER in fed and starved flies was still observed, but was much smaller in magnitude: 68% of fed flies and 96% of starved flies responded at the highest concentration (Figure 4F). This starvation-dependent enhancement of PER in bitter-silenced flies is likely to reflect the enhancement of the sugar-sensing pathway. These results reveal a state-dependent interaction between bitter and sugar neurons that affords a logic for the behavioral switch. In fed flies, bitter neurons strongly suppress the appetitive response to acetic acid mediated by sugar neurons. Hunger results in a behavioral switch that increases PER both by suppressing the bitter-sensing pathway and enhancing the sugar-sensing pathway.

### Acetic acid activates sugar- and bitter-sensing neurons

The observation that the appetitive response to acetic acid is mediated by sugar-sensing neurons whereas the aversive response is mediated by bitter neurons suggests that acetic acid is an unusual tastant capable of activating two opposing classes of sensory cells. We therefore performed two-photon imaging of taste sensory neurons to confirm whether acetic acid activates both sugar- and bitter-sensing cells. The genetically encoded calcium indicator *GCaMP6f* (Chen et al., 2013) was expressed in sugar-*(Gr64f-Gal4)* or bitter-*(Gr66a-Gal4)* sensing neurons. Imaging was performed on sensory axon termini in the subesophageal zone (SEZ) of the fly brain (Figure 5 – figure supplement 1). Strong GCaMP responses to acetic acid were observed in sugar neurons in both fed and starved flies, with the peak response to acetic acid about half of that observed with 500 mM sucrose (Figure 5A-D). We also examined the acetic acid response in sugar neurons of homozygous mutants carrying deletions in all eight sugar receptors *(ΔβGrs/ΔβGrs).* The response to sucrose in these mutants was reduced to the level of the response to water, whereas the response to acetic acid was not affected (Figure 5 – figure supplement 2). These results indicate that the response to acetic acid in sugar neurons does not require sugar receptors, a result consistent with the fact that sugar receptor mutants show strong PER to acetic acid (Figure 3D).

**Figure 5:**
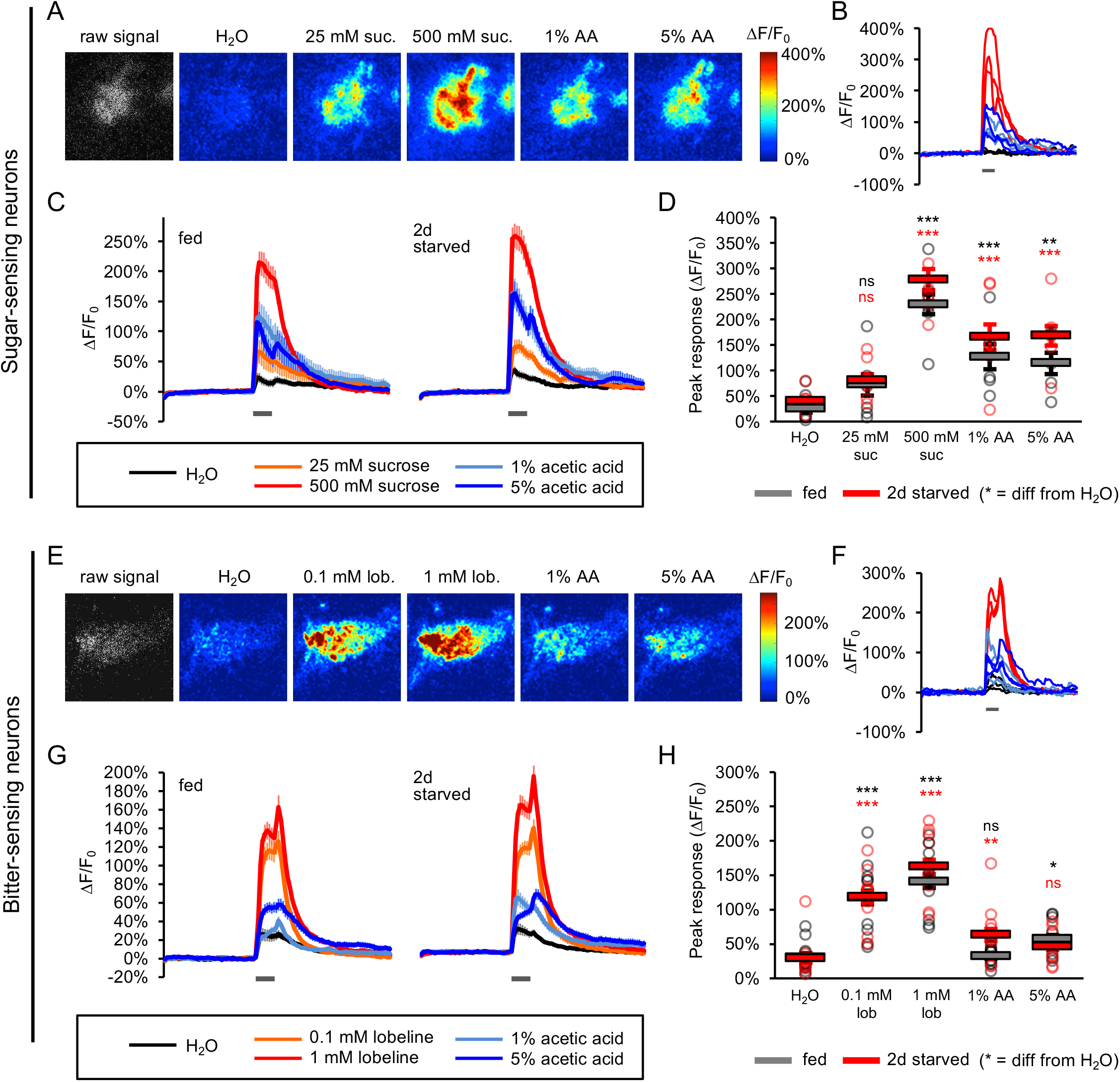
Acetic acid activates sugar- and bitter-sensing neurons. (A-H) Calcium imaging of taste sensory neurons reveals that acetic acid (AA) activates sugar-sensing (A-D) and bitter-sensing (E-H) neurons. (A, E) Spatial maps of GCaMP activation by each stimulus for individual flies. (B, F) Example ΔF/F_0_ traces for individual trials in the same flies shown in A and E, respectively. (C, G) Average GCaMP activation across all trials in all flies of each group. Grey bars indicate stimulus delivery (2 sec). (D, H) Peak response to each stimulus averaged across all trials for each group. Circles represent individual fly averages. Asterisks (color-coded by group) indicate responses that are significantly greater than the response of the same group to water (*p<0.05, **p<0.01, ***p<0.001, two-way ANOVA followed by Bonferroni post-tests). Peak sugar neuron responses of fed and starved flies (D) showed an overall difference (p<0.01, two-way ANOVA) but did not significantly differ for any individual tastant (p>0.05, all Bonferroni post-tests). Peak bitter neuron responses of fed and starved flies (H) did not significantly differ (p>0.05, two-way ANOVA). Data shown in this figure represent n=15-19 trials, 6 flies per group (sugar neurons) or n=30 trials, 10 flies per group (bitter neurons). We independently replicated these experiments with similar results (n=8 flies per group for both sugar and bitter neuron imaging; data not shown), but those data were not combined with the data shown here due to different imaging conditions.

Acetic acid also activated the bitter neurons with peak responses about 30% of those obtained with 1 mM lobeline (Figure 5E-H). The difference between the levels of activity elicited by lobeline and acetic acid may reflect different sensitivities of bitter neurons to the two compounds or the activation of a smaller subset of neurons by acetic acid. We therefore imaged the acetic acid response of four different subclasses of bitter neurons (Weiss et al., 2011) and observed that only one class exhibited a significant response when compared to the response to water (Figure 5 – figure supplement 3). This class of acetic acid-responsive neurons represents the S-b class, a finding consistent with a previous study demonstrating that the bitter cells in this class respond most strongly to acid tastants (Charlu et al., 2013). Thus, acetic acid activates both sugar- and bitter-sensing taste neurons.

We confirmed that acetic acid does not promiscuously activate all classes of taste neurons by imaging responses of water-sensing neurons, sensory cells in the labellum that respond to low osmolarity tastants (Cameron et al., 2010). Water-sensing neurons responded broadly to several taste stimuli at levels that were inversely related to their osmolarity (Figure 5 – figure supplement 4). When the low-osmolarity response of water-sensing neurons was blocked by adding the high molecular mass polymer polyethylene glycol (PEG) to each taste solution, acetic acid did not activate the water-sensing neurons beyond the level elicited by PEG alone (Figure 5 – figure supplement 4C-D). Acetic acid therefore activates water-sensing neurons solely by the osmolarity-sensing mechanism. Thus, the responses to acetic acid in sugar- and bitter-sensing neurons are specific and are likely to be mediated by receptors that recognize acetic acid.

We next compared the responses to acetic acid in sugar- and bitter-sensing neurons in fed and starved flies. In sugar neurons, starved flies showed a trend toward higher responses to 500 mM sucrose and both 1% and 5% acetic acid, but these differences did not reach significance (Figure 5C-D). Bitter neurons of fed and starved flies did not significantly differ in their responses to either lobeline or acetic acid (Figure 5G-H). The observation that only a subset of bitter neurons are activated by acetic acid may obscure hunger-dependent changes when measuring the response of the entire bitter neuron population. In addition, we observe substantial fly-to-fly variability in sensory responses that may also obscure small differences between fed and starved flies. We confirmed that the GCaMP6f-expressing flies used for imaging exhibit strong hunger-dependent increases in PER to both acetic acid and sucrose despite the weak hunger modulation of sensory responses (Figure 5 – figure supplement 5). Thus the effect of starvation on sensory neuron responses may be inadequate to explain the behavioral switch in the response to acetic acid. The striking effects of internal state on this behavior may therefore result from state-dependent modulation of both the sugar and bitter circuits downstream of the sensory neurons.

## DISCUSSION

Innate behaviors are observed in naive animals without prior learning or experience suggesting that they are mediated by neural circuits that are genetically determined. Meaningful stimuli such as the taste and smell of food elicit stereotyped behaviors that are observed in all individuals in a species. However, innate behavioral responses can also exhibit flexibility and can be modulated by experience, expectation, and internal state (Bargmann 2012; Ding and Perkel, 2014; Kim et al., 2017). Taste circuits in flies and mice appear to be anatomically and functionally programmed to elicit an innate appetitive or aversive response, ensuring that animals consume nutritious foods and avoid the ingestion of toxins (Liman et al., 2014). We have examined the response of *Drosophila* to acetic acid, a tastant that can be a metabolic resource but can also be toxic to the fly. Our data reveal that flies accommodate these conflicting attributes of acetic acid by virtue of a hunger-dependent switch in their behavioral response to this stimulus. Fed flies show taste aversion to acetic acid, likely a response to its potential toxicity, whereas starved flies show a robust appetitive response that may reflect their overriding need for calories. These opposing responses are mediated by two different classes of taste neurons. Acetic acid activates both the sugar and bitter pathways, which have opposing effects on feeding behavior. The choice of behaviors is determined by internal state: hunger shifts the response from aversion to attraction by enhancing the appetitive sugar pathway as well as suppressing the aversive bitter pathway (Figure 6).

**Figure 6:**
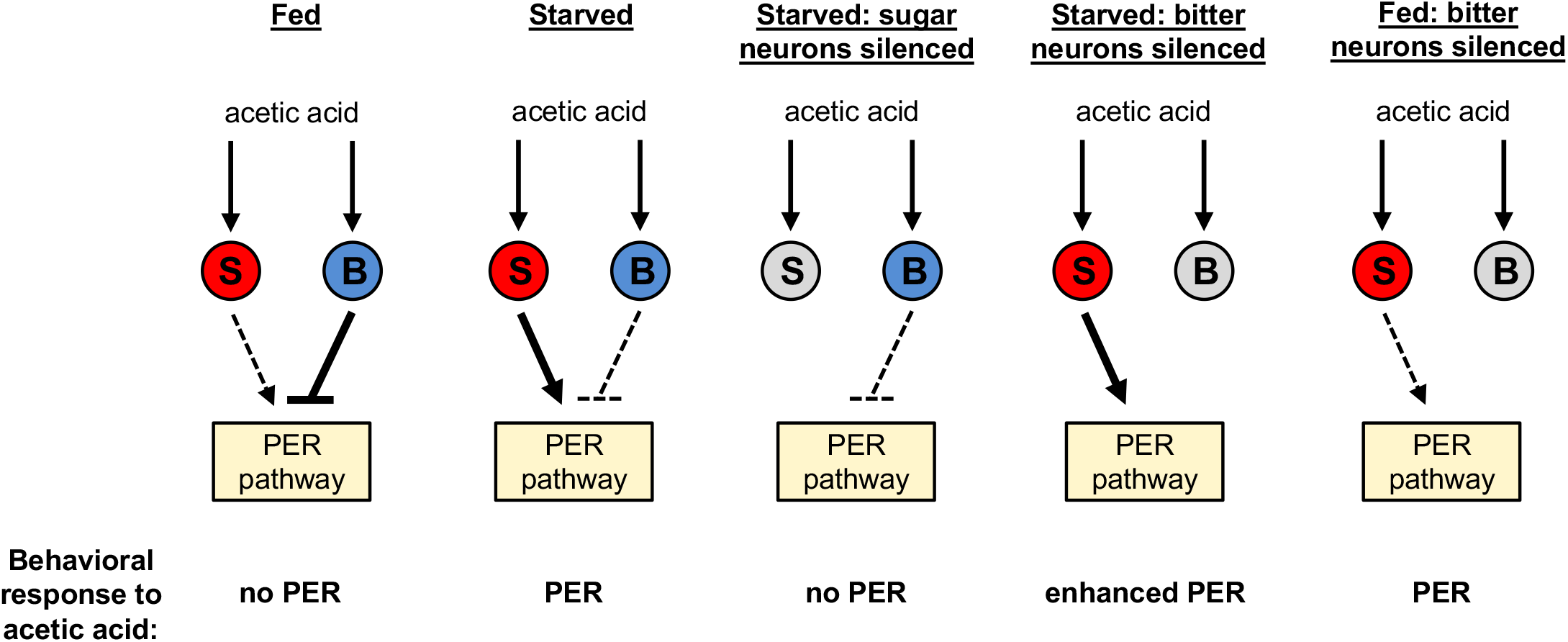
Model for a hunger-dependent switch in the behavioral response to acetic acid. Acetic acid activates both sugar- and bitter-sensing neurons (“S” and “B” respectively). Sugar-sensing neurons promote PER to acetic acid whereas bitter-sensing neurons suppress PER. The balance of these two pathways determines the behavioral response. Hunger both enhances the sugar pathway and suppresses the bitter pathway, most likely by acting downstream of sensory neurons. Thus the bitter pathway dominates in the fed state to suppress PER and elicit aversion, whereas the sugar pathway dominates in the starved state to elicit appetitive PER behavior. Silencing the sugar or bitter neurons (grey) shifts the balance of the two pathways to alter the behavioral response, as shown in the three right panels.

The adaptive response to acetic acid is dependent on two biological features, the ability of acetic acid to activate two classes of sensory neurons that elicit opposing behaviors and the state-dependent modulation of these two taste pathways. We observe that acetic acid activates both the bitter- and sugar-sensing neurons. The activation of bitter neurons by organic acids has been observed previously by electrophysiologic recording of labellar sensilla, and these acids result in taste aversion (Charlu et al., 2013). The observation that bitter neurons respond not only to organic acids but also to hydrochloric acid suggested that a subset of bitter neurons serve as a pH sensor eliciting taste aversion (Charlu et al., 2013). It remains unclear, however, whether acetic acid activation is mediated by this presumed pH-sensitive pathway or a distinct receptor mechanism.

We also observe that acetic acid activates sugar neurons, eliciting an appetitive taste response. This behavioral response is not observed upon exposure to low pH or acetate, suggesting the presence of a receptor on sugar neurons recognizing short chain aliphatic acids. A previous study demonstrated that whereas acids activate bitter neurons, acids inhibit rather than excite sugar neurons (Charlu et al., 2013). This result obtained by sensillar recordings contrasts with our observations that acetic acid activates sugar neuron axons. At present we have no explanation for these conflicting results. We observed acetic acid activation of sugar neurons with three different *Gal4* drivers dictating the expression of two different GCaMP variants (Figure 5A-D, Figure 5—figure supplement 2, and data not shown). The observation that acetic acid elicits PER in starved flies and this response is eliminated upon silencing of the sugar neurons is most consistent with the activation of sugar neurons by acetic acid.

The activation of different classes of sensory neurons by a single tastant has been observed for salt (Zhang et al., 2013), the long chain fatty acid, hexanoic acid (Ahn et al., 2017), as well as for acetic acid in this study. In both flies and mammals, low salt concentrations elicit attraction whereas high salt results in aversion and these opposing behaviors are mediated by distinct classes of sensory neurons (Zhang et al., 2013; Chandrashekar et al., 2010; Oka et al., 2013). Hexanoic acid, a caloric source, activates fatty acid receptors in sugar neurons and at high concentrations activates a different receptor in bitter neurons (Ahn et al., 2017). A logical pattern emerges in which tastants of potential value to the fly activate attractive taste pathways. These compounds may also be toxic and also activate aversive pathways either at higher concentrations or in different internal states. This affords the fly protection from the potential toxicity of excess, a protection that can be ignored under extreme conditions to assure survival.

Internal state can elicit profound behavioral changes that allow the organism to adapt to a changing internal world. Hunger, for example, results in enhanced food search and consumption, increased locomotion, changes in food preference, and altered olfactory and taste sensitivity (Sternson et al., 2013; Itskov and Ribeiro, 2013; Pool and Scott, 2014; Yang et al., 2015). Previous studies have observed that hunger enhances olfactory attraction to cider vinegar, increases sugar sensitivity, and decreases bitter sensitivity (Root et al., 2011; Inagaki et al., 2012; Inagaki et al., 2014). These effects of hunger represent gain changes; stimuli become more attractive or aversive. By contrast, our experiments reveal a qualitative change in the valence of acetic acid: hunger induces a switch from taste aversion to attraction.

Genetic silencing experiments indicate that this switch in starved flies results from inhibition of the aversive pathway, mediated by bitter neurons, and enhancement of the appetitive pathway mediated by sugar neurons. Conversely, in fed flies the aversive pathway is enhanced whereas the appetitive pathway is inhibited. Previous studies show that hunger modulates the behavioral response at least in part by modulating the activity of the initial neurons in this pathway, the sensory neurons (Inagaki et al., 2012; Inagaki et al., 2014). Our experiments imaging sensory neuron projections in the SEZ and those of Kain and Dahanukar (2015) fail to reveal significant differences in the response of either sugar or bitter neurons between fed and hungry flies. We did, however, observe a trend toward increased responses of sugar neurons to both sucrose and acetic acid in starved flies (Figure 5C-D).

Our data suggest that the striking state-dependent switch in the behavioral response to acetic acid likely reflects modulation of taste pathways downstream of the sensory neurons. The circuit from sensory neurons leading to proboscis extension remains largely uncharacterized but multiple nodes subject to modulation can be anticipated. The response to sugar extends to multiple behaviors beyond PER including ingestion, swallowing, and suppression of locomotion, each of which is likely to be modulated by hunger (Pool and Scott, 2014). Modulation at the level of sensory neurons affords gain control that will result in changes in all behaviors elicited by gustatory neurons. Modulation of downstream taste neurons facilitating sensorimotor transformations could afford a flexibility enabling independent control of different behavioral programs driven by the same taste stimulus. This affords genetically determined neural circuits mediating innate behaviors the opportunity for more complex modulation dependent on perception, motivation, and internal state.

## MATERIALS AND METHODS

### Fly stocks and maintenance

Flies were reared at 25°C and 70% relative humidity on standard cornmeal food. 2U was used as the wild-type control strain. All lines used for behavior were outcrossed into this background for at least 5 generations, with the exception of the octuple sugar receptor mutant *(ΔβGrs)* which contained too many mutations to outcross. PER assays were generally performed on 3-6 day-old mated females. Calcium imaging was performed on >1 week-old flies to ensure robust GCaMP6f expression, and PER assays for GCaMP6f-expressing flies were performed using flies of the same age.

All fly strains have been described previously: *Gr64f-Gal4* (Dahanukar et al., 2007); *Gr66a-Gal4* (Scott et al., 2001); *ppk28-Gal4* (Cameron et al., 2010); *Gr98d-Gal4, Gr22f-Gal4, Gr59c-Gal4,* and *Gr47a-Gal4* (Weiss et al., 2011); *UAS-Kir2.1* (Baines et al., 2001); *UAS-GCaMP6f* (Chen et al., 2013); *UAS-norpA-IR^31113^* (Masek and Keene, 2013); *poxn^ΔM22-B5^* and *poxn^ΔM22-B^ + SuperA* rescue (Boll and Noll, 2002); *Δ8Grs (R1, AGr5a;; ΔGr61a, ΔGr64a-f)* and *Δ8Grs* with transgenes for GCaMP imaging *(R1, ΔGr5a; Gr61a-Gal4, UAS-GCaMP6m; ΔGr61a, ΔGr64a-f)* (Yavuz et al., 2015).

### PER assay

Fed flies were taken directly from food vials for testing. Starved flies were starved with water (using a wet piece of Kimwipe) for the specified amount of time before testing. Flies were anesthetized on ice and immobilized on their backs with myristic acid. Unless otherwise specified, PER experiments were conducted by taste stimulation of the labellum. To ensure that we could deliver tastants to the labellum without contacting the legs, we immobilized the two anterior pairs of legs with myristic acid. For leg stimulation experiments (Figure 1E-F, Video 1, and Video 2), all legs remained free. Flies recovered from gluing for 30-60 min in a humidified chamber before testing.

Before testing PER, flies were water-satiated so that thirst would not affect their responses. PER to water (the negative control) was tested after water-satiation, followed by taste stimuli in ascending order of concentration. Flies were water-satiated again before each test. Each test consisted of two trials in which the solution was briefly applied to the labellum or legs using a small piece of Kimwipe. PER on at least one of the two trials was considered a positive response. Only full proboscis extensions, not partial extensions, were counted as PER. Flies were tested in groups of 15-20, and the percent of flies showing PER to each tastant was manually observed and recorded. Flies that did not respond to any taste stimuli were tested with 500 mM sucrose at the end of the assay. For experiments using only wild-type starved flies, which should always respond to high concentrations of sugar unless they are extremely unhealthy, flies that failed to respond to 500 mM sucrose were excluded from analysis. For experiments comparing fed and starved flies or starved controls and mutants, flies were only excluded from analysis if they appeared very sick.

For statistical analyses of PER, each group of 15-20 flies was considered to be a single data point. A minimum of three groups per genotype or condition were tested for each PER experiment. Because PER can vary substantially from day to day (likely due to changes in ambient temperature or humidity), control and experimental flies for the same experiment were always tested on the same days, and all experiments were repeated over at least three days.

To test directional PER, we contacted the left or right forelegs with acetic acid, alternating between sides every 1-2 trials. Flies were filmed and the videos were analyzed later. We only analyzed trials in which flies showed full PER to the stimulus. Flies often showed repeated extension to a single stimulation; at least one proboscis extension toward the left or right side was considered to be a lateralized response.

To test the role of olfaction, the third antennal segments and maxillary palps were removed with forceps while flies were anesthetized on ice. Surgery was performed prior to starvation, and after surgery flies were given ~30 min to recover in food vials before starvation. Control flies were anesthetized for the same duration as antennectomized flies.

### Calcium imaging

Flies for calcium imaging were taped on their backs to a piece of clear tape in an imaging chamber (see Figure 5—figure supplement 1). Fine strands of tape were used to restrain the legs, secure the head, and immobilize the proboscis in an extended position for tastant stimulation. A small hole was cut into the tape to expose the anterior surface of the fly’s head. A square hole along the anterior surface of the head was then cut through the cuticle, including removal of the antennae, to expose the anterior ventral aspect of the brain that encompasses the SEZ. The esophagus was cut in order to visualize the SEZ clearly. The dissection and imaging were performed in modified artificial hemolymph (Wang et al., 2003; Marella et al., 2006), which substitutes 15 mM ribose for sucrose and trehalose.

Calcium imaging experiments were performed using a two-photon laser scanning microscope (Ultima, Bruker) equipped with an ultra-fast Ti:S laser (Chameleon Vision, Coherent) that is modulated by pockel cells (Conoptics). Emitted photons were collected with a GaAsP photodiode detector (Hamamatsu) through a 60X water-immersion objective (Olympus). A single plane through the brightest area of axonal projections was chosen for imaging. Images were acquired at 925 nm at a resolution of 256 by 256 pixels and a scanning rate of 3-4 Hz.

Tastants were delivered to the labellum via a custom-built solenoid pinch valve system controlled by MATLAB software. Pinch valves were opened briefly (~10 ms) to create a small liquid drop at the end of a 5 μL glass capillary, positioned such that the drop would make contact with the labellum. Tastants were removed after a fixed duration by a vacuum line also controlled by a solenoid pinch valve. Proper taste delivery was monitored using a side-mounted camera (Veho VMS-004), which allowed for visualization of the fly and tastant capillary using the light from the imaging laser. At least three trials of each stimulus were given, with at least one minute rest between trials to avoid habituation.

Calcium imaging data were analyzed using custom MATLAB code based on the code used in Hattori et al. (2017). Images were registered within and across trials to correct for movement in the x-y plane using a sub-pixel registration algorithm (Guizar-Sicairos et al., 2008). Regions of interest (ROIs) were drawn manually around the area of axonal projections. Average pixel intensity within the ROI was calculated for each frame. The average signal for 20 frames preceding stimulus delivery was used as the baseline signal (F_0_), and the ΔF/F_0_ values for each frame were then calculated. The peak stimulus response was quantified as the average of the ΔF/F_0_ values for the three highest consecutive frames during tastant presentation.

### Statistical analyses

Statistical analyses were performed using GraphPad Prism, Version 4. Statistical tests and results are described in the figure legends. All graphs represent mean ± SEM. For *Gal4/UAS* experiments, statistical significance was attributed only to data points for which experimental flies that differed from both the *Gal4/+* and *UAS/+* controls in the same direction.

## ACKNOWLEDGMENTS

We thank Daisuke Hattori for MATLAB code and assistance with calcium imaging; Barbara Noro for advice and comments on the manuscript; Chris Rodgers and members of the Axel laboratory for helpful feedback and suggestions; Adriana Nemes, Phyllis Kisloff, Miriam Gutierrez, and Clayton Eccard for general laboratory and administrative support; and Hubert Amrein, John Carlson, Ulrike Heberlein, Kristin Scott, and the Bloomington Stock Center for providing fly strains. This work was supported by the Howard Hughes Medical Institute (R.A.) and the Simons Foundation (R.A.).

## AUTHOR CONTRIBUTIONS

A.V.D. and R.A. designed research; A.V.D., B.S., and A.Z. performed experiments and analyzed data; A.V.D. and R.A. wrote the paper.

---

**Figure 1 – figure supplement 1:**
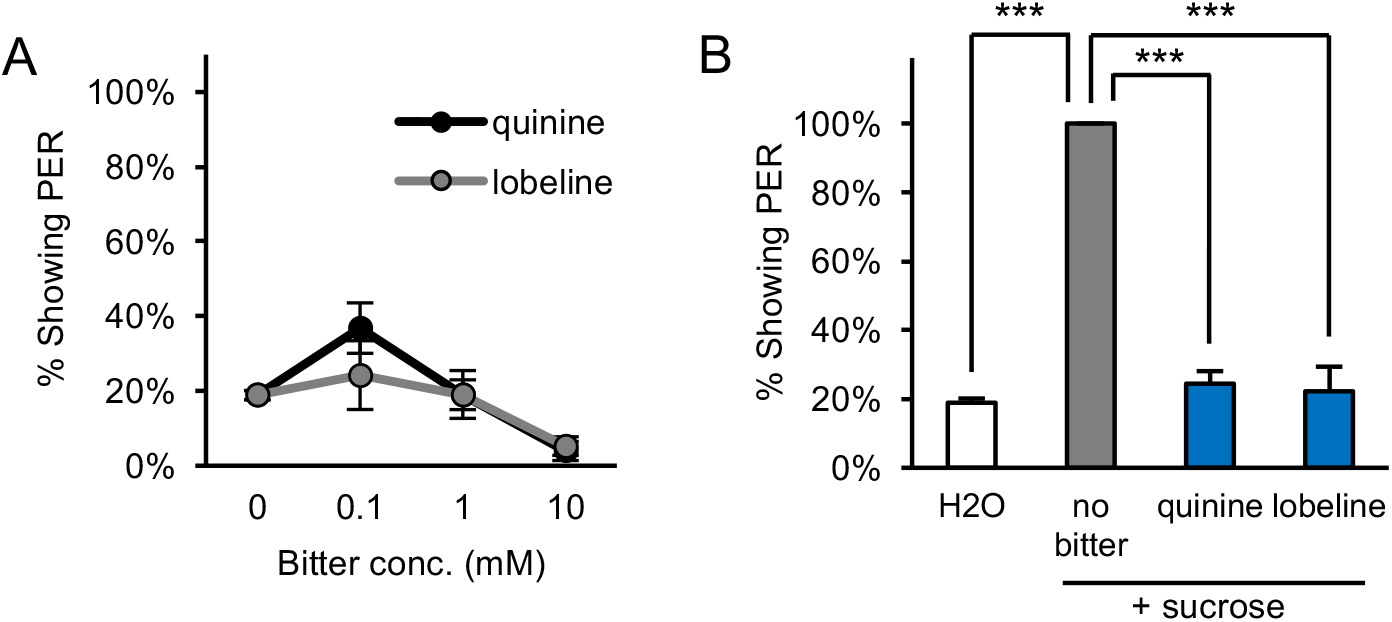
Starved flies show aversion to bitter compounds. (A) Two-day starved flies did not show consistent or strong PER to the bitter compounds quinine or lobeline (p>0.05 for lobeline, p<0.05 for quinine at 0.1 mM and 10 mM, one-way repeated measures ANOVA followed by Dunnett’s posttests comparing bitter compounds to water). (B) Two-day starved flies showed aversion to bitter compounds, as the addition of 10 mM quinine or 5 mM lobeline suppressed PER to 50 mM sucrose (***p<0.001, one-way repeated measures ANOVA followed by Bonferroni’s post-tests).

**Figure 1 – figure supplement 2:**
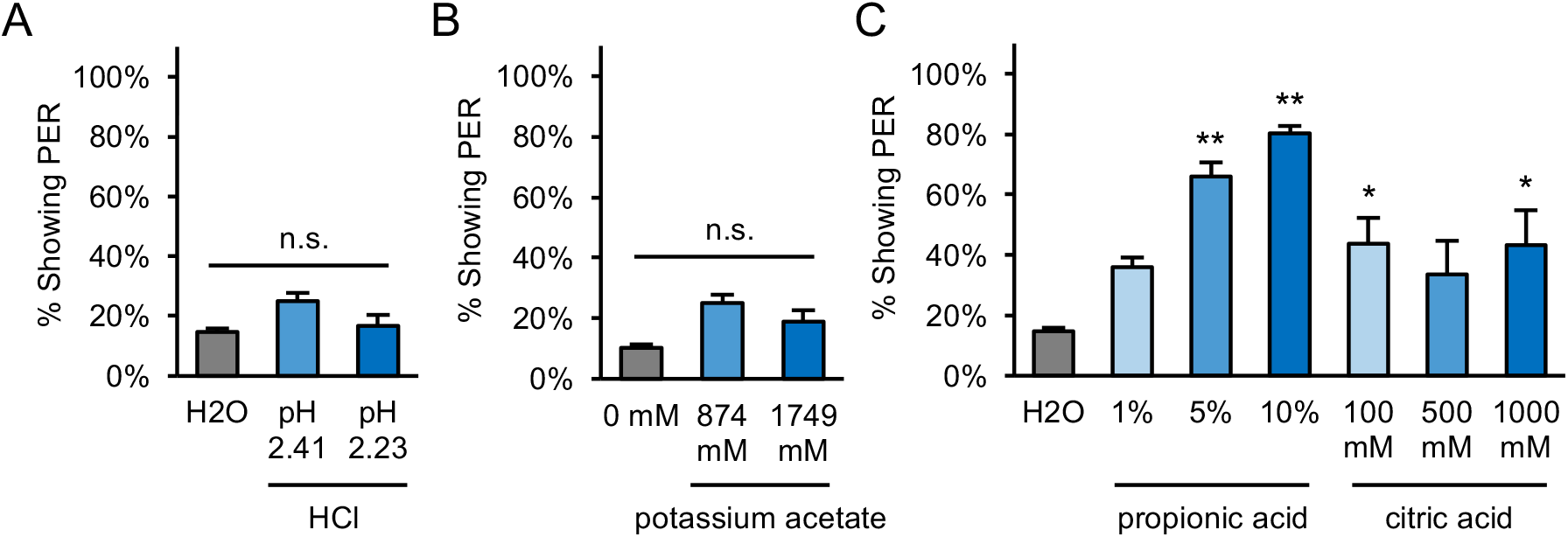
PER to other acids and acetate in starved flies. (A) Two-day starved flies did not show significant PER to hydrochloric acid (HCl) solutions prepared at the same pH values as measured for 5% and 10% acetic acid (pH 2.41 and 2.23 respectively). (B) Two-day starved flies did not show significant PER to potassium acetate presented at the same molarities as 5% and 10% acetic acid (874 and 1749 mM, respectively). (C) Two-day starved flies showed moderate to strong PER to propionic and citric acid. For all panels: *p<0.05, **p<0.01, one-way repeated measures ANOVA followed by Dunnett’s post-tests comparing each tastant to water.

**Figure 4 – figure supplement 1:**
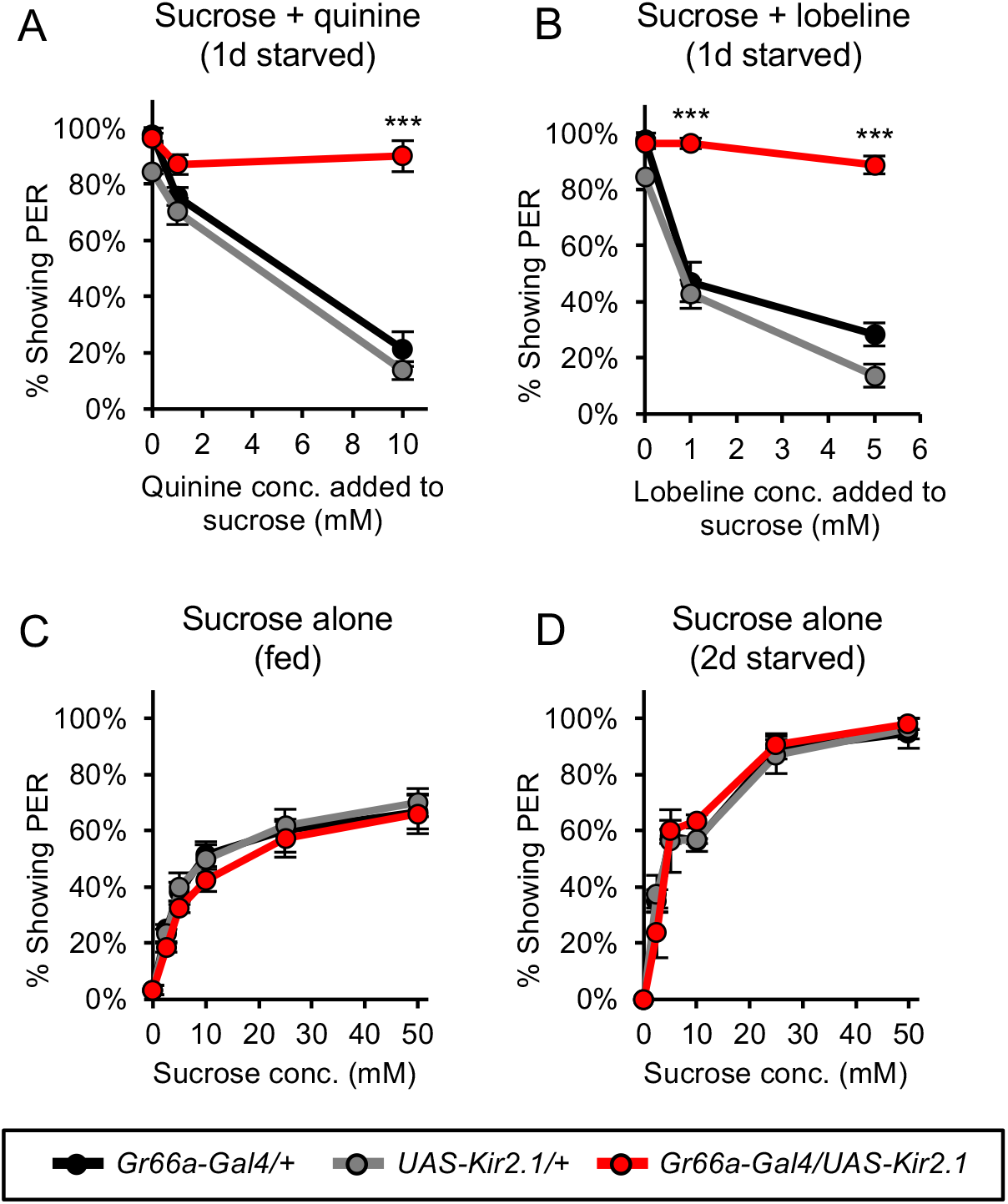
Silencing bitter-sensing neurons impairs bitter aversion but does not affect PER to sugar. (A-B) Silencing bitter-sensing neurons strongly reduced aversion to the bitter compounds quinine (A) and lobeline (B). Aversion was measured as the suppression of PER to 100 mM sucrose containing each of the bitter compounds. (C-D) Silencing bitter-sensing neurons did not affect PER to sucrose alone in fed flies (C) or two-day starved flies (D). For all panels: ***p<0.001, two-way repeated measures ANOVA followed by Bonferroni’s post-tests comparing experimental group to each control group.

**Figure 5 – figure supplement 1:**
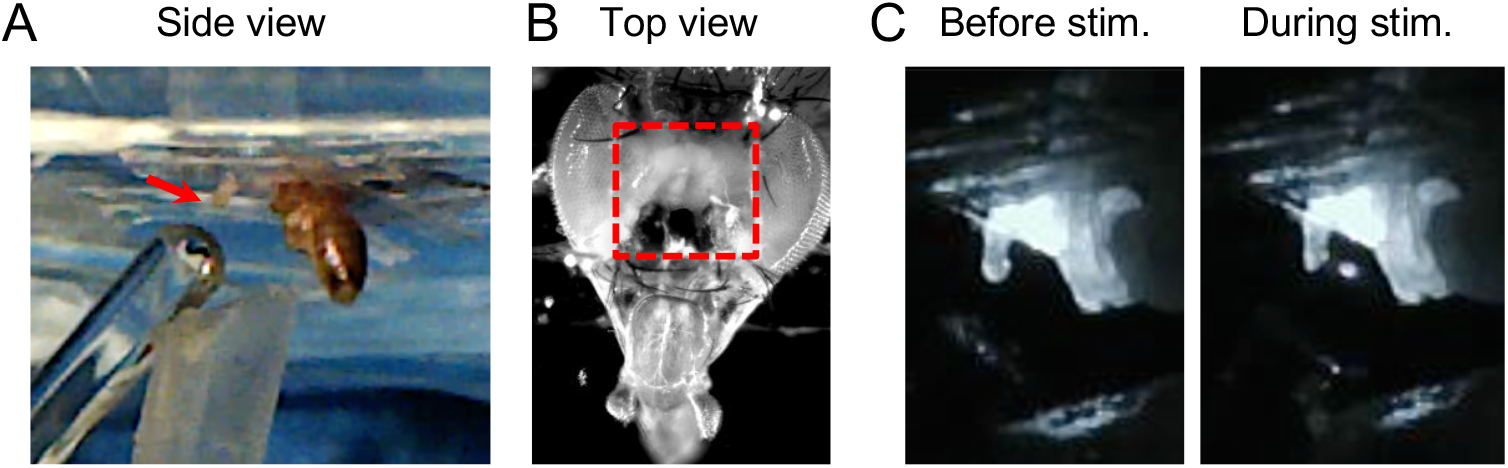
Calcium imaging setup for taste neuron imaging. (A) Side view of fly. Wings and legs were taped to allow unobstructed stimulation of the labellum (arrow). Tastant droplets were delivered to the labellum via a glass microcapillary and removed by a vacuum line, both of which were controlled by MATLAB software. (B) Top view of fly. Red box shows area of cuticle removed to expose the ventral brain, which includes the SEZ. (C) Still frames from video showing tastant delivery during imaging.

**Figure 5 – figure supplement 2:**
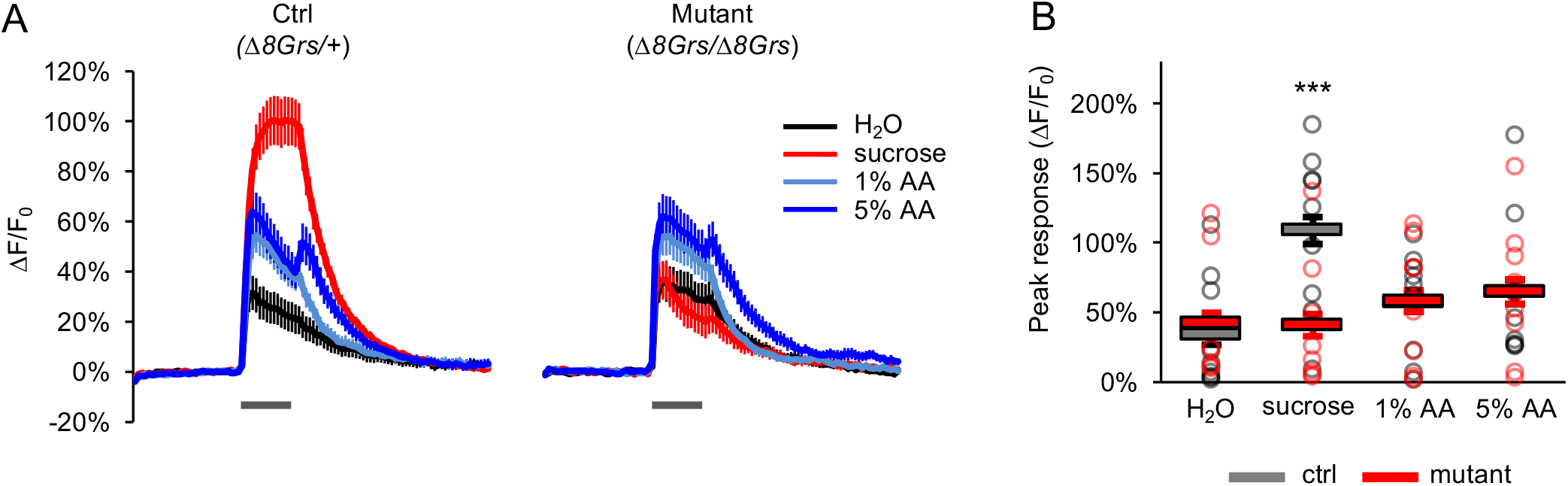
Acetic acid responses of sugar-sensing neurons in sugar receptor mutants are not affected. (A) Average GCaMP activation of sugar-sensing neurons in control flies (*Δ8Grs*/+) and mutant flies lacking all eight sugar Grs (*Δ8Grs/Δ8Grs*) in response to water, 100 mM sucrose, or acetic acid (AA). Both groups of flies were heterozygous for *Gr61a-Gal4* and *UAS-GCaMP6m* to enable expression of GCaMPΘm in sugar-sensing neurons. Grey bars indicate stimulus delivery (4 sec). (B) Peak response to each stimulus averaged across all trials for each genotype; circles represent individual fly averages. Only the response to sucrose differed significantly between genotypes (***p<0.001, two-way ANOVA followed by Bonferroni post-tests).

**Figure 5 – figure supplement 3:**
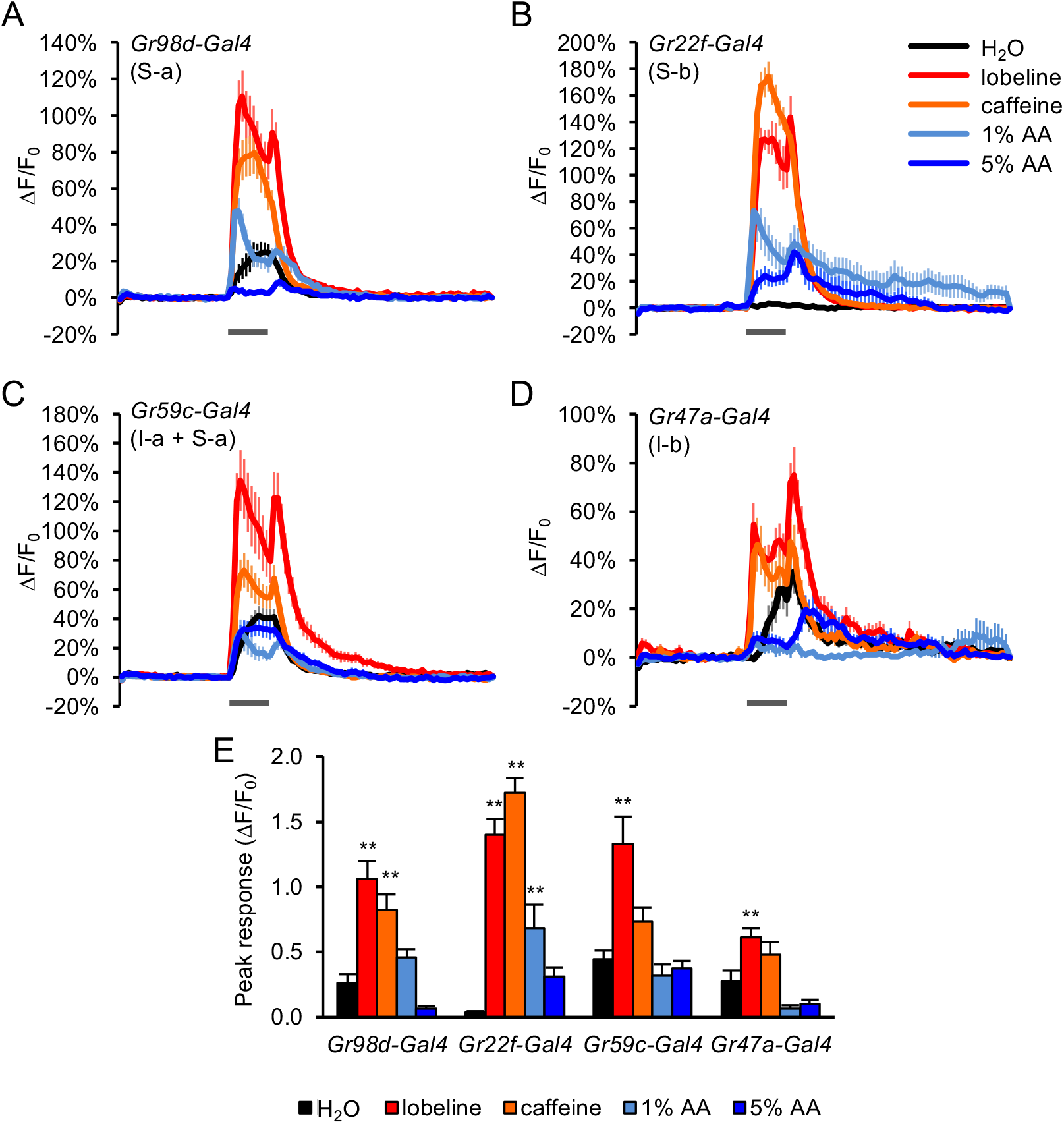
Acetic acid activates a subset of bitter-sensing neurons. (A-D) GCaMP responses of bitter neuron subsets were imaged using Ga/4 lines that label each of the four subclasses of bitter neurons (S-a, S-b, l-a, l-b) with complete or partial specificity, as noted in each panel. Flies were stimulated with water, bitter compounds (1 mM lobeline, 10 mM caffeine), and acetic acid (AA, 1% or 5%) for 3 sec (grey bars). Relatively weak GCaMP expression in neurons labeled by *Gr47a-Gal4* may account for their lower response magnitudes. (E) Peak GCaMP responses of each of the four bitter neuron subsets shown in panels A-D. Asterisks indicate responses that are significantly higher than the response to water (**p<0.01, one-way ANOVA followed by Dunnett’s post-tests). Only neurons labeled by *Gr22f-Gal4* showed a significant response to either concentration of acetic acid. Neurons labeled by *Gr98d-Gal4* did not show a significant difference in the peak response to acetic acid versus water, but the GCaMP traces for 1% acetic acid and water show clear differences in their timecourse (A), suggesting that these neurons may also respond to acetic acid.

**Figure 5 – figure supplement 4:**
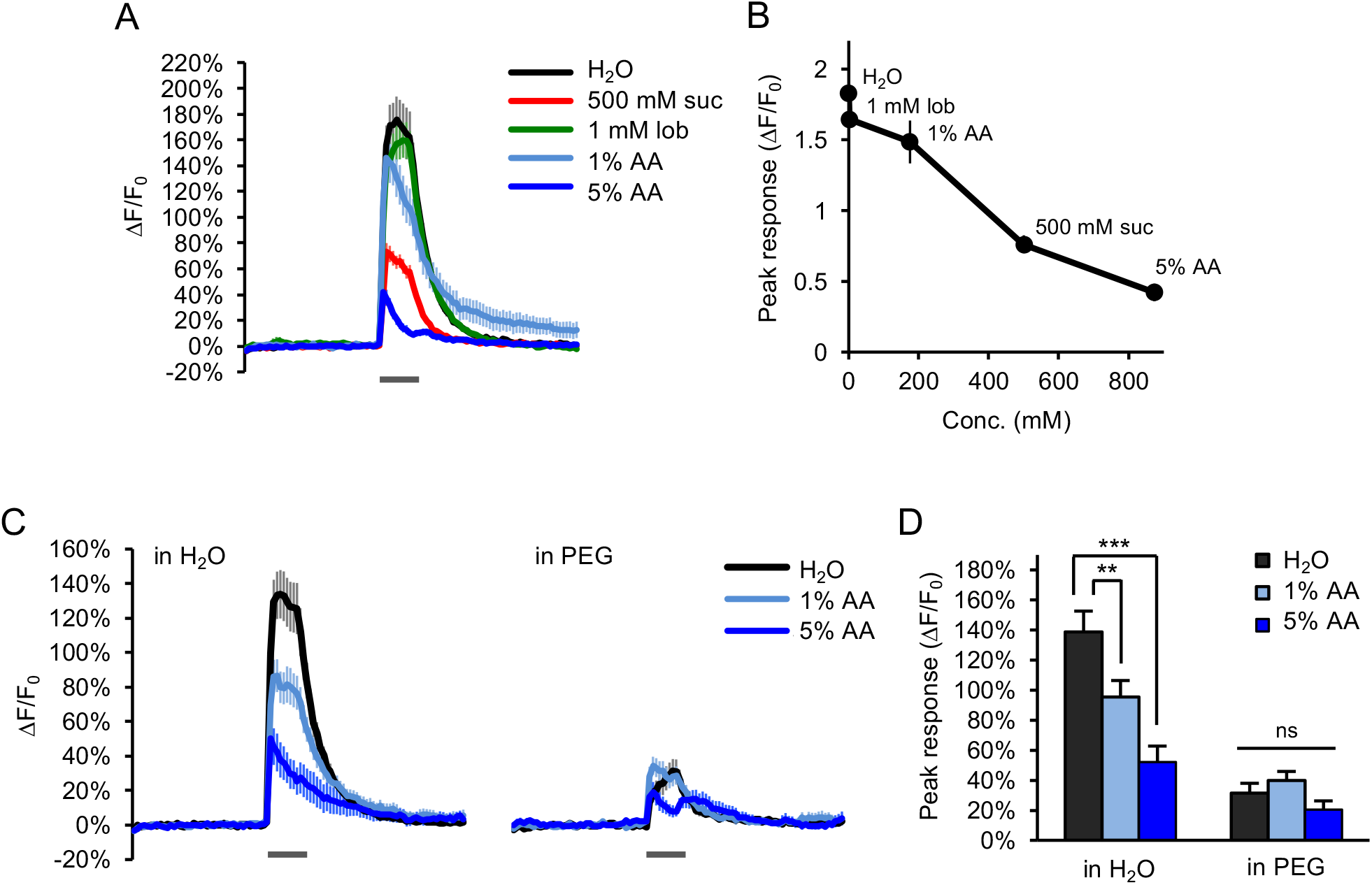
Water-sensing neurons are activated by acetic acid only in accordance with its osmolarity. (A-B) Water-sensing neurons labeled with *ppk28-Gal4* showed GCaMP responses to various taste stimuli (grey bar, 2 sec), including water, sucrose (suc), lobeline (lob), and acetic acid (AA). Responses decreased with increasing tastant molarity (B). (C-D) Adding PEG (3350 g/mol) to each tastant solution (10% wt/vol) strongly diminished the response to both water and acetic acid (p<0.001, two-way ANOVA). Responses to acetic acid in PEG were reduced to the same level as that of PEG alone (p>0.05, two-way ANOVA). **p<0.01, ***p<0.001, two-way ANOVA followed by Bonferroni’s post-tests.

**Figure 5 – figure supplement 5:**
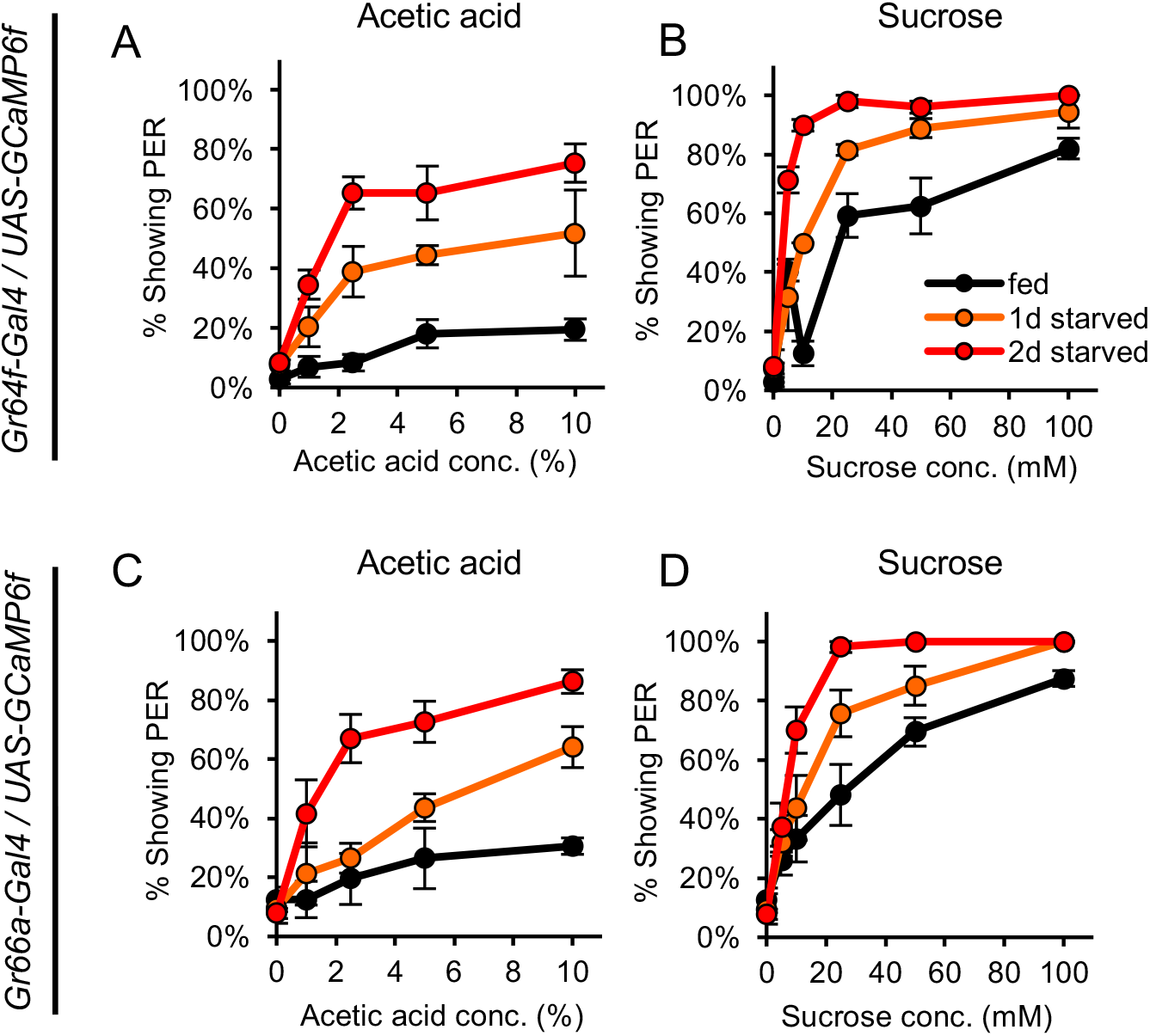
GCaMP-expressing flies show hunger-dependent changes in PER to acetic acid and sucrose. (A-D) Fed and starved flies expressing GCaMP6f in sugar-sensing neurons (A-B) or bitter-sensing neurons (C-D) were tested for PER to acetic acid (A, C) and sucrose (B, D). Both genotypes showed hunger-dependent changes in the level of PER to acetic acid (p<0.001 for *Gr64f-Gal4/UAS-GCaMPβf,* p<0.01 for *Gr66a-Gal4/UAS-GCaMP6f;* two-way repeated measures ANOVA) as well as sucrose (p<0.001 for both genotypes; two-way ANOVA).

